# Perception of Non-Verbal Prosody in Children with ASD

**DOI:** 10.1101/2020.12.07.414201

**Authors:** Aleksandra V. Maslennikova, Galina V. Portnova, Olga V. Martynova

## Abstract

Paralinguistic features of the speaker, such as prosody, temp, loudness, and dynamics, are an important marker of a person’s emotional state. The deficit of processing of emotional prosody could be preferably associated with the impairments in individuals with ASD’s social behavior. The following two groups of children participated in our study: 30 preschoolers from 4 to 6 years old in the target group (39.1 ± 6.4 scores by Childhood Autism Rating Scale), 24 preschoolers of the control group from 4 to 6 years in the control group. The prosody stimuli were the combination of syllables, said with intonations of “joy,” “angry,” “sadness,” “fear,” and “calm.” Fast Fourier transform (FFT) is used to analyze power spectrum density (PSD). The resulting normalized spectrum was integrated over unit width intervals in the range of interest (2 to 20 Hz with a step in 1Hz). Children with ASD, similarly to TD children, showed the most pronounced differenced of EEG in response to prosodics of fear and anger. The significant groups’ differences in PSD were detected for sad and joy intonations. Indexes of EEG differences between pleasure and painful intonations were significantly higher in the control group than children with ASD and between sadness and calm or joy and calm intonations. This paper makes up two main contributions: In general, we obtained that children with ASD have less response to a human voice’s emotional intonation. The physical characteristics of stimuli are more critical than a sign of emotions. The effect of EEG spectral power has hemisphere specialization in the healthy control group, but not in ASD children. Since spectral power for negative emotions in the target group is higher, we proposed that ASD children worse recognize positive emotions than negative emotions.

## 1. Introduction

Intonation or prosody of speech could be one of the most critical communication channels because it contains information about the emotional state of the speaker (Coutinho E, Dibben N, 2013). During a face-to-face conversation, the non-verbal information was transmitted and transduced to the brain through a hearing’s function using physical features such as phonation, pitch, loudness, and timbre encrypted in the speaker’s voice (Pernet, C. R., & Belin, P. (2012).). The non-verbal information transmission was evolutionarily more ancient than speech (Jespersen A.; Rubinshtein S., 1976; Linden F., 1981; Jakushin V., 1989), and the ontogenesis of non-verbal vocalization preceded the development of speech (Buck, R., 1975). Moreover, brain damage or anesthesia caused speech disorders led to impairment of primarily verbal speech and, secondly, to a deficit of non-verbal communication provided by more ancient brain structures and therefore were more resistant to a damaging effect. According to previous findings, the prosodic of speech could not depend on the meaning of the speech and often did not involve the speaker’s or the listener’s consciousness (Ramishvili D.I., 1978; Morosov V.P., 2011). In particular, during a conversation, intonation can be recognized as contradicting to the semantic component of speech (Eriksson, Anders; Francisco Lacerda, 2007; Liberman A, 2001).

The speech’s prosodic component showed that the emotional prosody of a speech depends on the harmonic characteristics of voice more than on other factors such as age, gender, profession, or meaning of the statement (Dmitrieva E.S. et al., 2009). Studies in psychoacoustics and experimental audiology oriented to the research of auditory transition also showed that for the same phrases with different intonations, there was a shift of the voice’s overtones, depending on the expressed emotion (Morosov V.P. et al., 1990). In particular, overtones of calm or moderate joy intonation were usually harmonious. In contrast, fear or anger prosodics had a disharmonious structure specific to each emotion and could be higher or lower than from harmonic overtones. The described effect, called “the quasiharmonic phenomenon,” underlined the forming of a speaker’s psychological portrait by listener based on non-verbal features of the speaker’s voice (Degottex, 2014; Singh, J. B., & Lehana, P. K., 2019). Thus, the audio signal’s frequencies were the primary source of information about the speaker’s emotional state.

The ability to understand the emotional context underlay social perception and the capacity for communication. Recent studies (Hellbernd et al., 2018) demonstrated that auditory non-verbal information was dominant in processing social information and prevailed over the visual during perception of the interlocutor’s emotional states. At the same time, the deficit of emotional auditory perception was one of the main symptoms in patients’ emotional sphere with schizophrenia and individuals with autistic spectrum disorder (Tobe, 2016; Bonfils KA, 2019).

The impaired recognition of emotional prosody is most consistent with the clinical picture of the autism spectrum disorder (ASD) associated with a deficit of social perception (Le Gall, 2018). It could be detected even in adults with high-functioning autism (Globerson E., 2015), who not only perceived worse but also reproduced worse basic emotions in speech (Hubbard DJ, 2017). The causes of impaired recognition of emotional prosody in children with ASD are still not clear enough. However, several researchers suggested that impaired perception of affective prosody in children with ASD could also be associated with dysfunction of brain regions involved in speech prosody’s perception. In particular, previous findings demonstrated that individuals with ASD recognized affective prosody worse than the control group due to impaired connectivity of the amygdala and anterior cingulate cortex and the superior temporal regions (Rosenblau, 2017). Schelinski S. (2016) also indicated the reduced activation of these brain areas in subjects with ASD during human voice perception. Other findings revealed that perception of verbal prosody was associated with a function of lower frontal and motor brain areas in children (Correlia Al. et al., 2019) and the right frontal regions (Ghacibeh GA et al., 2003; Bateman JR, 2019). Finally, the resent findings indicated that deficit affecting the processing of emotional prosody cannot be accounted for merely in terms of impairments in one isolated brain area (Philip, 2010) but could be preferably associated with a wide range of brain functions contributes to the impairments in the social behavior of individuals with ASD.

In this study, we hypothesized that impaired recognition of emotional prosody in children with ASD could be associated with their preference for prosody’s physical characteristics during perception, such as harmonious or disharmonious structure, pitch, and loudness. Moreover, we assumed that difficulties in learning and processing social-related aspects of prosody contributed to subtle communication in interpersonal relationships and problems in understanding emotions.

## 2. Materials and Methods

### 2.1. Participants

The following groups of children participated in our study: 30 preschoolers from 4 to 6 years old (Mean = 5.2; SD = 0.8), 20 boys and ten girls with ASD (39.1 ± 6.4 scores by Childhood Autism Rating Scale (CARS), Second Edition) and 24 preschoolers of the control group from 4 to 6 years old (Mean = 5.4; SD = 0.9), 17 boys and 7 girls (18.2 scores by CARS). The criteria that contributed most to exclusion were the presence of comorbid neurological or psychiatric conditions, a history of moderate to severe brain injury any epileptic activity on EEG. Children of both groups did not take any medication for at least two months before the study. All the subjects (or their authorized relatives) signed a written informed consent before participating in the study. The Ethics Committee approved the study’s protocol of the Institute of Higher Nervous Activity and Neurophysiology of RAS.

### 2.2. Stimuli

The prosody stimuli consisted of combination of syllables “mi-me-ma-mo-mu-my”, said by a professional actress with intonations of “joy”, “angry”, “sadness”, “fear” and “calm”. The sounds had duration 1500 ms and loudness 70 dB and had following characteristics: wave file type, channel format-stereo, sampling frequency-48 kHz, 24 bits. The detailed physical characteristics of stimuli were depicted on Figure 1A.

**Figure 1.**
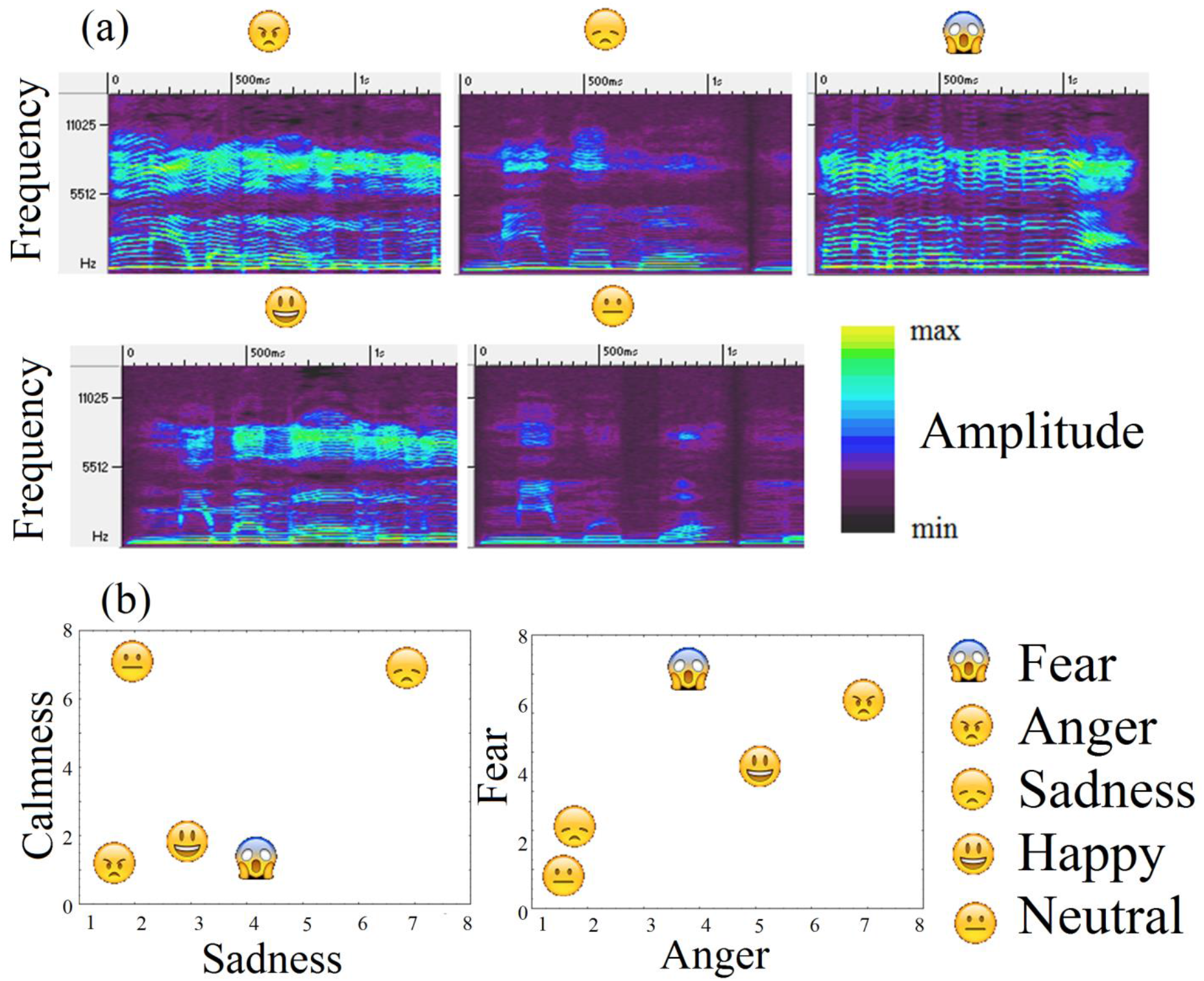
Stimuli characteristics (A): pitch and volume for all types of intonations and their assessments (B).

**Figure 2.**
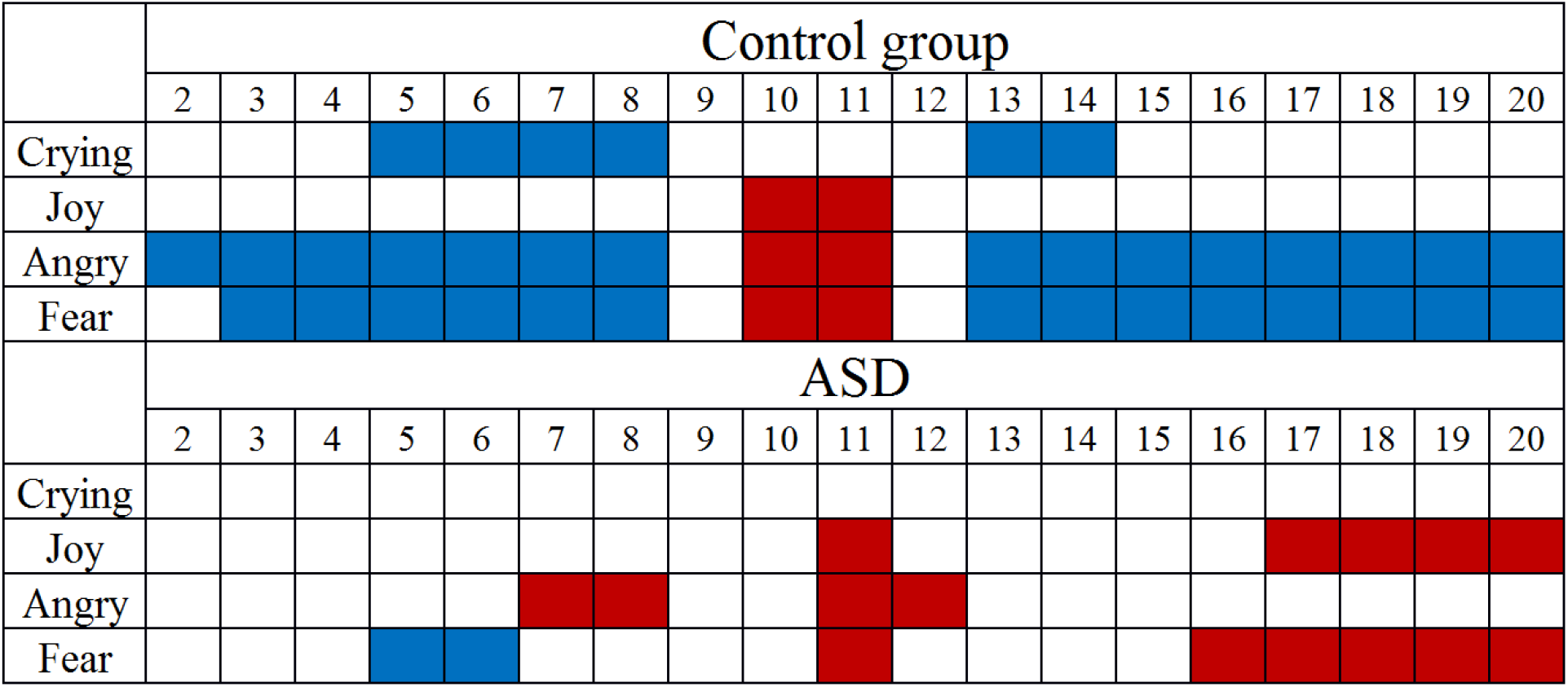
PSD differences between calm and all types of intonations. Significant increase of PSD marked red, decrease–blue for all frequency steps (2 to 20 Hz in step of 1 Hz).

### 2.3. Stimuli assessments

We asked children to assess stimuli using analog scales: we asked children to mark the desired emotion using scales Calmness (0–9), Sadness (0–9), Anger (0–9), and Fear (0–9). Unfortunately, due to children’s difficulties in understanding and following the instructions, only 21 from 24 children in the control group and 8 from 30 children with ASD could cope with this task. The results of the assessment of stimuli by children of the control group were depicted in Figure 1B.

### 2.4. Procedure

Stimuli were presented through the speakers with constant and comfortable subject volume in a randomized sequence through a randomized duration of inter-stimulus interval (from 1000 to 3000 ms). Children were asked not to move and sit with eyes opened.

### 2.5. EEG registration

Resting-state EEG was acquired using a 19-channel EEG amplifier, “Encephalan,” with the recording of polygraphic channels (Poly4, Medicom MTD, Taganrog, Russian Federation) for 10 minutes. A sampling rate was 250 Hz. The amplifier bandpass filter was nominally set to 0.05–70 Hz. AgCl electrodes (Fp1, Fp2, F7, F3, Fz, F4, F8, T3, C3, Cz, C4, T4, T5, P3, Pz, P4, T6, O1, and O2) were placed according to the International 10–20 system. The electrodes placed on the left and right mastoids served as joint references under unipolar montage. The vertical EOG was recorded with AgCl cup electrodes placed 1 cm above and below the left eye, and the horizontal EOG was acquired by electrodes placed 1 cm lateral from the outer canthi of both eyes. The electrode impedances were kept below 10 kΩ.

### 2.6. EEG preprocessing

Continuous EEG corresponding to stimulation and each subject’s resting state was cleaned from eye movements by an ICA-based algorithm in the EEGLAB plugin for MatLab 7.11.0 (Mathwork Inc.). Muscle artifacts were cut out through manual data inspection. The continuous resting-state EEG of each subject was filtered with a band-pass filter of 0.5–30 Hz. The final duration of each analyzed EEG fragment was 178 ± 22.3 s.

### 2.7. Data analysis

#### 2.7.1. Power spectral density (PSD)

Fast Fourier transform was used to analyze PSD. The EEG spectrum was estimated for every 178 ± 22.3 s intervals. The resulting normalized spectra were integrated over intervals of unit width in the range of interest (2–3 Hz, 3–4 Hz, 19–20 Hz).

#### 2.7.2. EEG spaces

We also used method of EEG spaces which was previously successfully applied on children with ASD (Portnova, 2018) to visualize how close/distant the prosodic sounds and the resting state were according to EEG data. The main steps of the procedure were following:

EEG of each stimuli and resting state fragments was divided into small non-overlapping epochs of 8 seconds (approx. 10-12 pieces).

FFT (absolute value) was calculated for the epochs in 2-20 Hz band for the subset of the electrodes (F3, F4, F7, F8, T3, T4, T5, T6, P3, P4, C3, C4). Fp1 and Fp2 were excluded as they don’t usually contain any useful information about cognitive processes, O1 and O2 were excluded as they cover visual cortex but the presented stimuli are auditory, Fz, Cz and Pz were excluded as the reflection of emotions on EEG is asymmetric.

The distance between each pair of the stimuli was calculated: for each frequency bin (with 1/16 Hz step) two samples of FFT values (of the epochs of these fragments) were compared using Mann-Whitney U-test (threshold p <0.05). The distance was equal to the percentage of differing frequency bins. Emotional stimuli and background states were placed onto a plane using multidimensional scaling method, namely Sammon projection. Each stimulus was depicted as a emoticons (Figure 1). So, the distances between the stimuli types on the plane were as similar as possible to the distances calculated by FFT values. This similarity was always good enough to claim the projection is legit.

The resulting pictures (obtained for each subject) have arbitrary rotation because of the Sammon projection algorithm and different sizes because of EEG’s high individuality. Before the averaging over the group, these pictures should be standardized. We used scaling to equalize the size (the sum of squared distances to the circles/squares from the ‘center of mass’) and rotation/reflection so that fearful prosody was on the top of the picture and the background state with eyes opened and calm prosody were on the left and right sides correspondingly laying on a horizontal line.

After standardization individual pictures were averaged over groups. So, these pictures show relational distances between emotional sounds based on how much the corresponding EEG data differ in terms of rhythms magnitudes. The program to conduct these calculations was implemented on C# programming language by the lab’s engineer mentioned in Acknowledgements section.

### 2.8. Statistical analysis

Linear and repeated measures ANOVAs with following post-hoc comparison (Bonferroni, p <0.05) was used to determine group effects on EEG metrics and stimuli assessments. We analyzed a possible association of the EEG metrics with the ratings of subjective evaluation of emotional triggers and age using Spearman correlation analysis corrected for multiple comparisons by cluster-based permutation test using clustering method (Matlab toolbox for BCI) with 500 permutations at each node (the Bonferroni corrected p-value of 0.05); the clusters of differences for the EEG metrics were also calculated by cluster-based permutation test using clustering method using Wilcoxon signed-rank test. The group differences in behavioral assessment results were calculated using the Mann-Whitney test. Only significant (p <0.05) correlations and differences were reported.

## 3. Results

### 3.1. Power spectral density (PSD) Emotional Prosody compared to calm

#### 3.1.1. Sad Prosody

Children with ASD didn’t have significant differences between sad and calm intonations. In contrast, the control group children showed a decrease of delta and theta-rhythm PSD in occipital areas and a decrease of the beta-rhythm 12–14 Hz (F(2, 104)=7.984, p=,0067) for sad prosody compared to calm.

#### 3.1.2. Joy prosody

Children with ASD showed increase of alpha-rhythm’s PSD (11–12 Hz) for joy prosody compared to calm in central and parietal areas. Similar increase of alpha-rhythm was also detected in TD children (F(1, 52)=6,825, p=,0093): the PSD of 10–12 Hz alpha-rhythm increased in parietal areas for joy prosody compared to calm. At the same time, this prosody induced also significant increase of the beta-rhythm PSD (17–20 Hz) in occipital areas in children with ASD that was not detected with TD children (F(1, 52)=5,817, p=,0110).

#### 3.1.3. Angry Prosody

The angry prosody compared to calm induced significant increase of alpha1-rhythm PSD (7–9 Hz) in the left temporal areas and alpha2-rhythm (11–13 Hz) PSD in central area in children with ASD.

During listening to angry prosody compared to calm children of control group showed decrease of lower frequency PSD (2–8 Hz) in temporal, central and parietal areas, increase of alpha-rhythm PSD (10-12 Hz) in central and parietal areas and decrease of beta-rhythm PSD (13-20 Hz) in frontal areas.

#### 3.1.4. Fear prosody

Children with ASD showed decrease of theta-rhythm’s PSD (5–7 Hz) for fear prosody compared to calm in central and temporal areas, increase of alpha-rhythm PSD (10–12 Hz) and beta-rhythm (13– 20 Hz) in central areas.

Children of control group also showed similar to children with ASD significant decrease of low frequency band’s PSD (3–7 Hz) for fear prosody compared to calm (F(1, 52)=7,2 p=,0035) and increase of alpha-rhythm PSD (10–12 Hz) (F(1, 52)=6,82 p=,0089), but also (unlike to children with ASD) decrease of the beta-rhythm (13–20 Hz) PSD in frontal areas (F(2, 104)=8.642, p=,0008).

Thus, children with ASD similarly to TD children showed the most pronounced differenced of EEG in response to prosodics of fear and angry, which were also similar and different only in opposite beta-rhythm changes.

### 3.2. Differences between joy and sad prosodics (Permutation clustering test using Wilcoxon Matched Pairs Test)

The most pronounced group differences were detected for sad and joy intonations. Moreover, the results of permutation clustering analysis revealed significantly higher low-frequency band’s (2– 8 Hz) PSD (z=3.24, p=0.003) and beta band’s (16–20 Hz) PSD (z=3.27, p=0.001) for joy compared to sad prosodic only in the control group (Figure 3). In children with ASD, these differences didn’t pass Bonferroni correction.

**Figure 3.**
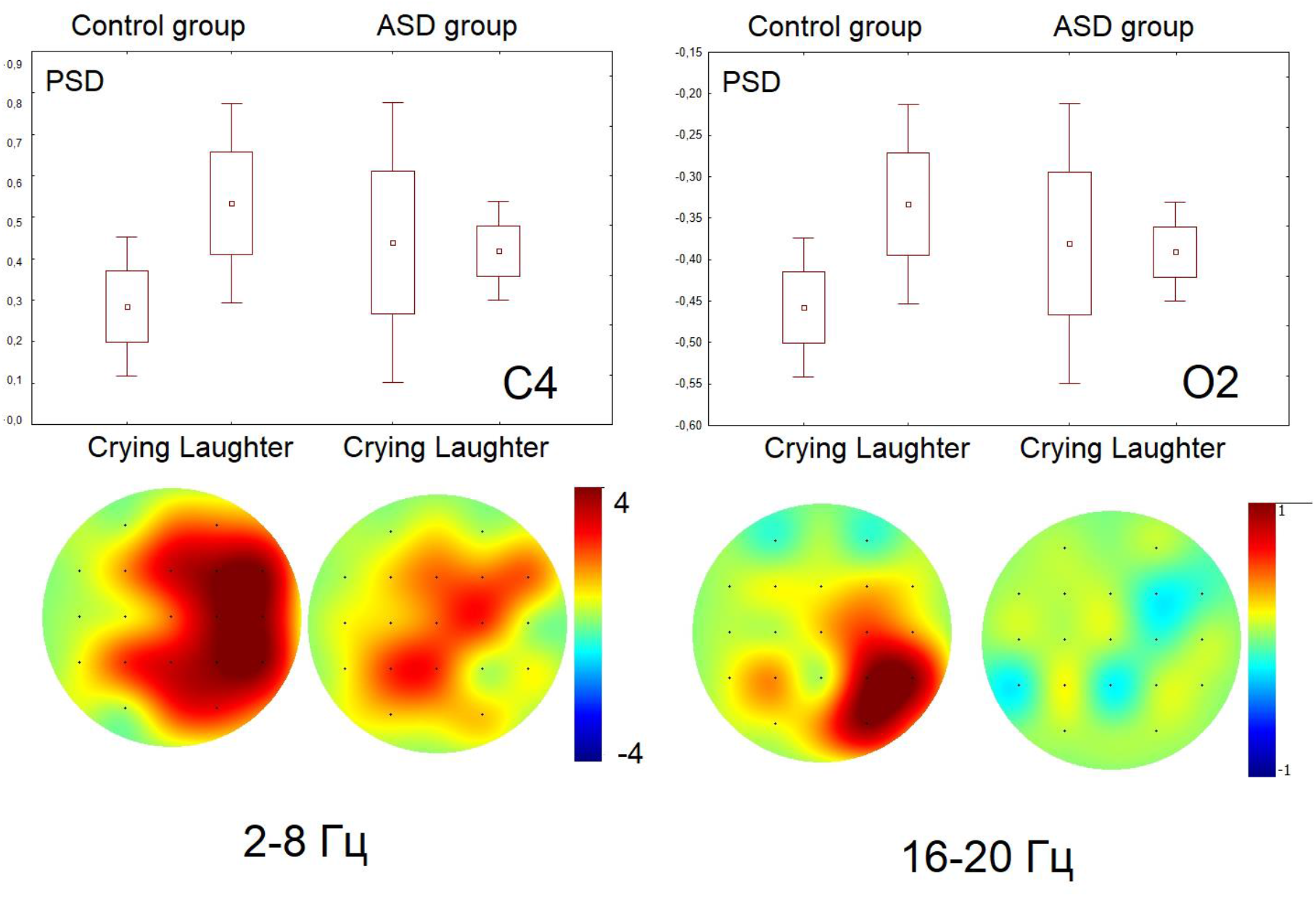
Topography of PSD coefficient between group healthy control and ASD in 2–8 Hz span for sad (crying) and joy (laugher) prosody.

**Figure 4.**
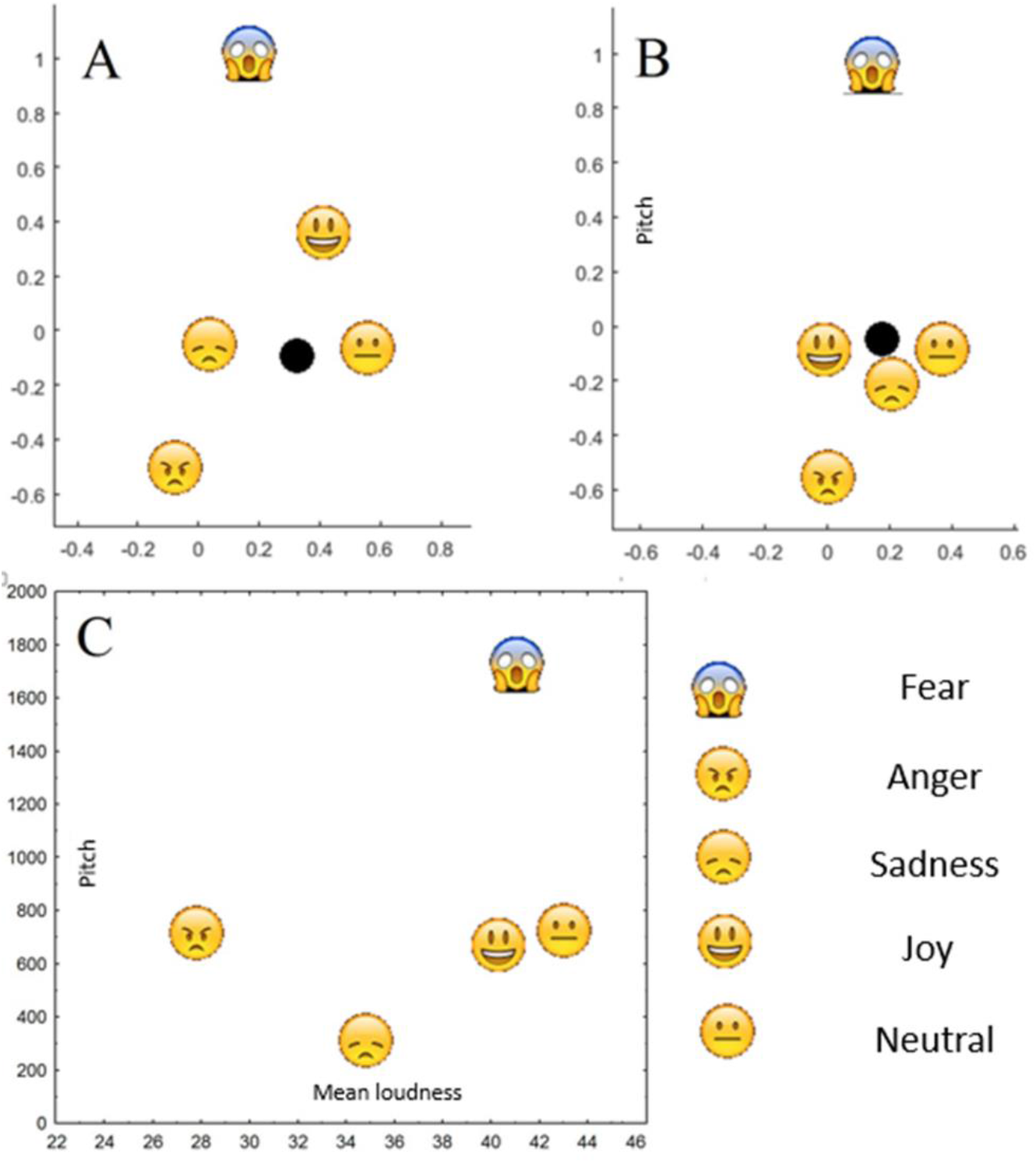
Space of EEG indexes differences for all type of intonations. (A) EEG spaces of children of control group. (B) EEG spaces of children with ASD. Black dot–resting state EEG.

### 3.3. EEG spaces with indexes

Indexes of EEG differences between joy and sad intonations were significantly higher in the control group compared to children with ASD (z=2.98, p=0.006), as well as between sadness and calm (z=2.92, p=0.008) and joy and calm intonations (z=3.16, p=0.001). Other group differences between EEG indexes were not detected.

### 3.4. Correlation analysis between EEG indexes and physical parameters and assessment of sounds

The significant correlation between EEG indexes and differences of prosodics’ pitches were detected in both groups of subjects (r=0,68, p=0.002). At the same time, the difference in the assessment of these sounds by scale “Sadness” significantly correlated with the EEG index of difference between these prosodic only in TD children (r=0.64, p=0.001) and was not detected in children with ASD even on-trend (p=0.89). Based on the higher difference of EEG response between joy and sad and calm prosodics in the group of TD children, the difference in emotional stimuli processing could be suspected.

## 4. Discussion

1986; Shriberg et al., 2001; Rapin and Dunn, 2003; Paul et al., 2005). Horie M, et al. l. (2017) studied preschool and early school children’s ability in comparison with the control group and children with attention deficit hyperactivity disorder to reproduce and recognize emotions in a speech on a large sample. He found that children with ASD are the worst at recognizing unpleasant emotions, while children with ADHD do a little better. The data of electrophysiological studies confirm this: the amplitude of the P300 component of the negativity of the mismatch of the evoked potential for words pronounced in different intonations is lower in children with ASD than in healthy subjects (Lindstrom R, 2016, 2018; Charpentier J, 2018). According to Krestar ML et al. (2019), in the task of identifying the intonations of semantically calm words, the subjects more often guessed a sad intonation rather than joyful or calm.

Although data regarding brain stem involvement are scarce, studies using cortical evoked potentials in patients with the autism spectrum (especially those with Asperger’s syndrome) have demonstrated insufficient speech coding. They have linked this deficit to poor receptive prosody. For example, adults with Asperger’s disorder who listened to a woman’s name, spoken in a calm manner or with a dismissive, sad, or commanding affect, had relative difficulty in identifying emotional connotations compared to controls and also showed significant differences in the negativity of mismatch (MMN, response reflecting encoding acoustic changes), including longer delays, lower amplitudes, and fewer evoked responses (Kujala et al., 2005). In a second study (Korpilahti et al., 2006), boys with Asperger Syndrome were presented with a woman’s name at two different fundamental frequencies to express affection or command affection. Their N1 responses (reflecting the onset of the stimulus) were delayed and decreased in amplitude compared to controls, and their MMN responses were earlier, larger, and had atypical laterality. The most recent study using MMN in this population showed an increase in the response (amplitude) in individuals with ASD with constant discrimination between the signs of both the main and vowel stimuli. In contrast, this effect disappeared when the condition included deciphering phonemes with changes in pitch (Lepisto et al., 2007). These findings are similar to Lepisto and colleagues’ previous work, indicating that adults and children with Asperger’s disorder (Lepisto et al., 2006) and children with autism (Lepisto et al., 2005) had an increased MMN response to sounds, which deviates in height from the standard stimulus. In this study, both standard and deviant stimuli had a constant pitch throughout the sound. However, they also showed a decrease in P3a responses (orientation response) to changes in pitch in speech, albeit not variant, in syllables.

The study of stem potentials for words spoken with different intonations has shown that some children with ASD demonstrate pitch tracking deficiencies (Russo N. et al., 2008). A significant influence of the coefficient of verbal intelligence in the control group on the latency of the evoked potential’s average components was detected during emotional stimuli perception. In contrast, the value of non-verbal intelligence was positively correlated with a decrease in the latency of the P300 component.

Topographic differences in rhythm distribution showed hemispheric specialization for intonation of joy: neurotypical children showed higher activation in alpha-rhythm (10-12 Hz) band in the right parietal-temporal lobe, ASD children, in contra, in the same region and frequency band in left. Considering that PSD differences between groups were found in the central and parietal regions of both hemispheres, which were previously associated with the processing of speech intonation, we can conclude that children with ASD are worse at recognizing the emotional context of speech.

For other intonations, all changes in power were associated with the left hemisphere and the central areas of both hemispheres. Besides the fact parietal lobes of the right are related to the processing of speech intonation, we suggested that children with ASD are worse at recognizing the emotional context of speech. The differences between laughing and crying in the control group have lateralization, which indicates the involvement of emotionally significant processes. The right hemisphere is associated with analyzing emotional processes, which is especially important when processing speech intonation (Schirmer, A., & Kotz, 2006).

In the control group children, an increase in power was noted precisely in the right area for the intonation of joy. In contrast, in children with ASD, lateralization was not revealed. This data may indicate a decrease in sensitivity to the intonation of joy in speech, difficulties in its recognition. The change in the power of slow-wave activity in the control group prevailed in the occipital region over a more comprehensive frequency range. Simultaneously, similar changes of PSD in children with ASD were detected only at a frequency of 4-5 Hz in the frontal lobe (Zietsch, B. P. et al., 2007). At the same time, previous studies demonstrated that an increase in the power of slow-wave activity is characteristic of patients with depression (Fernández, A. et al., 2005), as well as in subjects with sleep deprivation (Cajochen, C., Foy, R., & Dijk, DJ, 1999).

## 5. Conclusions

This paper makes up several main contributions. In general, we obtained that children with ASD have less power of response to the emotional intonation of a human voice. The physical characteristics of stimuli are more critical than the color of emotions. The effect of EEG spectral power has hemisphere specialization in the control group, but not in ASD children. Since spectral power for negative emotions in the target group is higher, we proposed that ASD children worse recognize positive emotions than negative.

## Author Contributions

Data curation, G.V.P. and O.V.M.; Formal analysis, G.V.P.; Investigation, A.M.; Writing– original draft, A.M.; Writing–review & editing, O.V.M.

**Publisher’s Note:** MDPI stays neutral with regard to jurisdictional claims in published maps and institutional affiliations.

